# Phylesystem: a git-based data store for community curated phylogenetic estimates

**DOI:** 10.1101/013862

**Authors:** Emily Jane McTavish, Cody E. Hinchliff, James F. Allman, Joseph W. Brown, Karen A. Cranston, Mark T. Holder, Jonathan A. Rees, Stephen A. Smith

## Abstract

**Motivation:** Phylogenetic estimates from published studies can be archived using general platforms like Dryad (Vision, 2010) or TreeBASE (Sanderson *et al*., 1994). Such services fulfill a crucial role in ensuring transparency and reproducibility in phylogenetic research. However, digital tree data files often require some editing (e.g. rerooting) to improve the accuracy and reusability of the phylogenetic statements. Furthermore, establishing the mapping between tip labels used in a tree and taxa in a single common taxonomy dramatically improves the ability of other researchers to reuse phylogenetic estimates. Because the process of curating a published phylogenetic estimate is not error-free, retaining a full record of the provenance of edits to a tree is crucial for openness, allowing editors to receive credit for their work, and making errors introduced during curation easier to correct.

**Results:** Here we report the development of software infrastructure to support the open curation of phylogenetic data by the community of biologists. The backend of the system provides an interface for the standard database operations of creating, reading, updating, and deleting records by making commits to a git repository. The record of the history of edits to a tree is preserved by git’s version control features. Hosting this data store on GitHub (2014) provides open access to the data store using tools familiar to many developers. We have deployed a server running the “phylesystem-api”, which wraps the interactions with git and GitHub. The Open Tree of Life project has also developed and deployed a JavaScript application that uses the phylesystem-api and other web services to enable input and curation of published phylogenetic statements.

**Availability:** Source code for the web service layer is available at https://github.com/OpenTreeOfLife/phylesystem-api. The data store can be cloned from: https://github.com/OpenTreeOfLife/phylesystem. A web application that uses the phylesystem web services is deployed at https://tree.opentreeoflife.org/curator. Code for that tool is available from https://github.com/OpenTreeOfLife/opentree

## 5 Introduction

Characterizing and systematizing relationships among species has been a goal of biologists since Linnaeus (1758). The phylogenetic systematics “revolution” of the 1960’s focused most of the effort toward this goal on the task of estimating phylogenetic relationships. The rapid growth in availability of molecular data, the development of models and software implementations for inferring phylogenies, and the appreciation of the explanatory power of phylogenetically-aware comparative methods (e.g. Felsenstein, 1985) have led to dramatic increases in the number of published phylogenetic hypotheses.

Unfortunately, capturing the inputs and outputs of a phylogenetic analysis in a rich, standardized form is difficult and error prone. There is no single standard for phylogenetic data, and developers of phylogenetic software often create new formats or extend existing formats in ways that make them incompatible with other programs. Some archiving services, such as Dryad (Vision, 2010), have reacted to this challenge by accepting a wide range of inputs and making few or no guarantees to users of the database that the records will be in any particular form. This encourages sharing of data by making the submission process fast and easy. However, if the authors submitting the data are not conscientious in their explanations of the data, it can be difficult for users of the data to reliably extract all of the necessary information from the archive, or the data may not contain sufficient metadata for reuse.

On the opposite end of the spectrum, some databases require data providers to conform to strict standards with respect to input format and content. Examples of this approach are TreeBASE (Sanderson *et al*., 1994) and its successor TreeBASE version 2 (Piel *et al*., 2009). Since 1994, TreeBASE databases has served as the primary archival data store for phylogenetic estimates and the data that these trees are based on. As of 2014, TreeBASE contains 4076 studies and over 12,800 trees (TreeBASE, 2014). Many journals encourage or require that each publication that estimates phylogenetic trees cite a TreeBASE deposit containing the associated trees. TreeBASE produces a much more feature-rich interface for phylogenetic queries than Dryad, because each TreeBASE submission is parsed thoroughly and converted to a set of records in a relational database. The downside of this approach is that it requires that submissions correspond to a uniform format. Without constant updating, that format may not reflect new types of phylogenetic data being generated.

Unfortunately, tree estimation tools read and write file formats which are idiosyncratic and often so terse that they omit useful information about the analysis. See Stoltzfus *et al*. (2012) and Cranston *et al*. (2014) for discussions of this topic and other challenges relating to the archiving of phylogenetic estimates. As a result, the TreeBASE submission process usually requires the authors of a data package to reformat their data, correct the rooting of the tree, alter the labels of the tips of the tree, *etc*. Because these steps are often taken after the analyses for a publication are complete, it is likely that errors introduced in the process of preparing a submission will not be corrected until an observant user tries to reuse the data. If there were a rich, universally used file format for phylogenetics analyses, then import constraints such as those used by TreeBASE would be less of a hurdle, because the critical metadata regarding rooting, labels, *etc*. would be stored alongside the tree itself. The XML-based NexML specification (Vos *et al*., 2012) defines such a format, but it is not currently used widely by tree estimation tools.

Thus, the current state of phyloinformatic archival infrastructure is ripe for improvement in numerous ways. One such opportunity is the development of a data store that allows the community of biologists to improve the accuracy and re-usability of published phylogenetic statements. We describe here an implementation of one such system, which we call “phylesystem” because it uses a set of versioned text files on the filesystem. The goal of phylesystem is not to replace systems like Dryad or TreeBASE, but to complement them by storing phylogenetic statements and associated metadata in a consistent format while retaining the history of edits that were made to the data themselves. We anticipate that most of the trees stored in phylesystem will be associated with permanent, static archives elsewhere (usually either TreeBASE or Dryad).

### 5.1 Approach

One of the motivations of the design of phylesystem was to support the Open Tree of Life’s need for a curated set of rooted trees that have been aligned to a common taxonomy. The Open Tree of Life project strives to support infrastructure for phylogenetic research in an open way that encourages the community of biologists to participate in the collection and curation of our knowledge of the phylogenetic relationships of life on Earth. Thus, our goals when designing phylesystem were to build:

1. A data store capable of:

a. Storing trees from thousands of published studies. We do not intend to store the data upon which the phylogenetic estimates were based.
b. Storing a few exemplar trees from each study. We do not anticipate that the system will be used to store thousands of trees from an individual study (such as each sampled tree from a bootstrap or MCMC analysis).
c. Handling rich annotation of the trees, the taxa to which they refer to, and metadata describing the analysis that produced the trees.
d. Storing the full history of changes made to a study and its trees, including an identifier to indicate who made the changes.
2. Web services around the data store to support a user-friendly curation application run in the user’s browser. These include services for validation of the files against a published schema.
3. A loosely coupled system that would allow the community of biologists to interact with the data in a wide variety of ways.

From these requirements, we chose to implement phylesystem as a set of software wrappers around a data store which consists of a git (Git, 2014) repository of phylogenetic statements serialized as a JavaScript Object Notation (JSON) (Crockford, 2006). The data model used for the JSON is a close derivative of the NeXML standard for data interoperability (Vos *et al*., 2012), and can be converted back and forth using the peyotl library (https://github.com/OpenTreeOfLife/peyotl; manuscript in preparation).

Our expectations that the system will need to store limited amounts of data for each study (points #1a and 1b above), and that most studies will require relatively few edits, imply that raw performance of the basic database operations is unlikely to be a bottleneck for most uses of phylesystem. This made it feasible to use text files as the format of the data store. Thus far, in the six months since deployment of the study curation interface, the mean number of edits per study is only 2.7. The total size of the stored data (in JSON format with a line ending after every field) is only approximately 150MB (not counting the git database that stores the edit history).

The requirement (#1d, above) of maintaining a history of changes (including an identifier for the editor) fits naturally with software version control systems (VCSs). We note that wiki engines such as Gollum use git version control as a backend database, and the idea of using git as general database has been discussed by others (*e.g*. Keepers, 2012).

The requirement (#1c above) to support rich annotations is met by the NeXML data standard. This standard was designed to allow arbitrarily complex annotations of the entities that are crucial to phylogenetic statements. Because phylesystem does not store character data, the entities included are the operational taxonomic units (OTUs), trees, nodes, and edges.

We opted to represent NeXML as JSON due to the widespread use of JSON and to make it efficient for the server to provide data for a client-side curation application that is written in JavaScript (requirement #2 above). We developed some syntactic conventions for converting XML to a terse JSON representation. This JSON is described below as otNexSON. The adoption of these conventions reduced the web service data payload size by approximately 50% relative to naive representation of NeXML in JSON. This made it more feasible to load each study into the memory of the client’s browser. Despite the departure from the NeXML syntax, phylesystem maintains the ability to export and ingest NeXML files. Fundamentally the data model of the system is the data model of NeXML.

The final requirement (#3) of producing a maximally open, loosely coupled system is consistent with the use of a distributed VCS. Distributed VCSs do not require a single, “primary” repository. Rather, each copy of the repository maintains a copy of the entire history, and sharing documents between repositories is a peer-to-peer interaction.

The copy of the phylesystem repository that is listed under the Open Tree of Life organization on the GitHub website can be considered the canonical version of the data store. Indeed, this is likely to be the most easily accessible and highest profile clone of the phylesystem repository. Nevertheless, other users of git are free to fork that repo and maintain their own variants of the data store. Any such clone of the phylesystem repository maintains the ability to pull in changes from biologists who contribute edits to the Open Tree of Life organization clone (for example, via Open Tree of Life web applications).

The Open Tree of Life project makes available all of the code for phylesystem under permissive, BSD-style licenses, and the project makes no claim of ownership to any of the data in the repository. For new files deposited through the web application, the interface strongly encourages application of a CC0 copyright waiver, allowing for later deposit in Dryad. Other data, including those files that originate from TreeBASE, do not have an explicit data license (although we believe that copyright does not apply to any of the data; see Patterson *et al*., 2014).

In most database-driven web services, the group of core maintainers who have shell access and database permissions are the only people who have full access to the data in a data store. Posting frequent dumps of a database can allow other users to obtain local copies of the data for expensive, *ad hoc* calculations. However, a local version of the data is clearly a second-class instance and the maintainers of the canonical version of the database have *de facto* ownership of the project. We hope that adopting a truly distributed backend for the data store will reinforce the goals of the Open Tree of Life project to build tools that can be used and controlled by the entire community of biologists (rather than a few claiming ownership on the data store in perpetuity).

Currently the phylesystem web services use GitHub authentication for “curators” who edit studies. The requirement that editors authenticate allows the provenance of each edit to include a user name for the curator. The use of GitHub credentials allows the Open Tree of Life project to circumvent the need to maintain a database of users and passwords (or hashes). Thus, there is no private database that would be required for someone to fork and maintain their own version of the corpus (on GitHub or elsewhere). The only private information relevant to phylesystem that is kept by the Open Tree of Life project are the ssh-private keys that allow the project’s web servers to push updated data to the GitHub-hosted clone of the repository. In the future pull requests from forks of the phylesystem repo will be able to contribute data to the main repository. In order to maintain data quality, the phylesystem API (typically via the curation webapp) is currently the only method to add studies to the database.

## 6 Methods

The basic workflow of a typical curation session is depicted in Fig. 1. The steps involved are:

1. The user loads the curation webpage in his/her browser. This fetches the JavaScript curator application into the web browser.
2. A list of studies from phylesystem is loaded from information in a study indexing service (called “OTI”, see below).
3. The user selects the study he/she would like to view.
4. The curator application requests the study in otNexSON form from the phylesystem server.
5. The curator application also receives the git commit SHA (a checksum of the content) of the repository at this point.
6. The user may browse the study and download representations of the data in NeXML, otNexSON, NEXUS Maddison *et al*. (1997), or Newick formats without logging in.
7. If the user wants to edit a study, he/she must have a GitHub account (which are available free of charge) and must authenticate at this point.
8. The user may correct tip labels and map OTUs to a taxonomy with the help of taxonomic name resolution services (the “taxomachine” web services, described at https://github.com/OpenTreeOfLife/taxomachine. Tip labels are mapped to the comprehensive Open Tree Taxonomy, OTT (https://github.com/OpenTreeOfLife/opentree/wiki/Open-Tree-Taxonomy manuscript in preparation), so that all trees in the data store can be directly compared. Users can also fix the rooting information about the tree or edit the metadata. The study data is modified in the memory space of the browser and maintained there until the curator chooses to save the study.
9. To save, the curator application uses an HTTP request to the API and includes the data (in otNexSON format), the starting SHA, and a commit message to support sensible history.
10. The phylesystem-api code validates the otNexSON (rejecting the request if the data do not conform to a legal otNexSON document).
11. If the edited study is a legal otNexSON document, then a new git commit is created with the SHA provided in the API request as the parent commit. The commit is placed on a “work-in-progress” branch in the git history to assure that the data are stored with no chance of conflict.
12. If the study in question has not changed in the master branch since the parent SHA, then the edit can safely be merged to the master branch. If this is the case, the merge is done and the work-in-progress branch is deleted. If the version of the study on the master branch *has* changed (e.g. if two users are simultaneously editing the same study), then the merge is not done. The data will be saved on the server, but not merged into the master branch. If the commit is not automatically merged, then the merge must be completed manually by a curator at a later time, as there is the potential for conflicts between changes made by different simultaneous edits.
13. If the new edit was successfully merged to the master branch, then an event is triggered to tell the server to push the new master branch changes to the GitHub version of the repository.
14. A response is returned to the curation application indicating whether or not the edited study was merged to the master branch. This response includes the new SHA that will serve as the parent for a future commit.
15. If the push event was triggered in step #12, the updated master branch will be pushed to GitHub.
16. Using GitHub webhooks as a callback mechanism, the push generates a POST that triggers reindexing of the affected study by the OTI tool.

**Figure 1.**
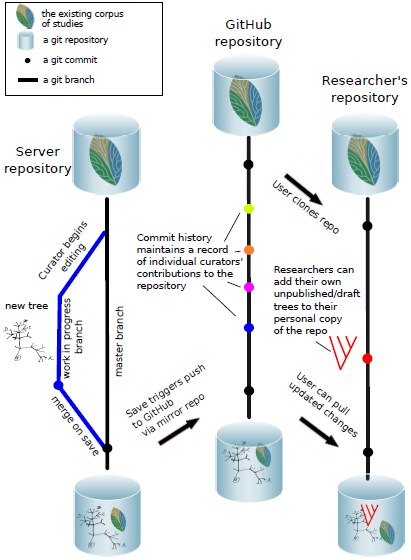
Workflow diagram demonstrating the process triggered by a curator editing a study using the curator app. Upon saving, a work in progress branch is generated on the server repository. If the study edited has not changed on the server, the work in progress branch is merged to master. Merges to master trigger pushes of commits, via the server mirror repository, to the Open Tree of Life GitHub repository. Curation of different studies can proceed independently, and on save each will be merged to master and trigger a push to GitHub. The record of curation changes is readily accessible (here represented with different colors for each curator). Users can easily access changes to the data store, and pull those changes into their own copy of the repository.

## 7 Implementation

Currently phylesystem is implemented in a python application using web2py for the web framework. Most of the functionality for validating the data and creating the git commits is accomplished using parts of the peyotl library. An alternative implementation of the web services has also been written in the Pyramid web framework.

### Thread safety

In the current implementation, the creation of a new commit and merge are done with git’s “checkout”, “add”, and “commit” commands. This means that the repository must use a mutual exclusion (mutex) lock for the duration of these events to ensure that a single thread completes this series of operations. Failing to lock the repository would make it possible for the HEAD reference (which will define the parent of a commit) to be moved as the result of a separate transaction. This would result in the git history not correctly reflecting which version of the study was displayed to the user by the curator application. Because of the mutex lock, the system can assure that the user’s edits appear as direct descendants of the repository version that provided the data to the curator application when the editing session began (step 4 above).

If the system encounters heavier use, then the wrappers around git may need to be modified to use lower level git commands. There are available commands that allow one to construct a commit without checking out the parent commit’s version of all files. These enhancements would limit or eliminate the need to lock the repository.

### Sharding

The phylesystem software supports having the data spread across multiple git repositories, which we refer to as “shards.” Currently, all of the studies fit within one git repository, so one shard suffices. Sharding can help avoid hitting limits on git repository size (for GitHub hosting) and can reduce contention for the mutex lock mentioned above.

The shard repositories contain the otNexSON files and a few files that are used to create a unique study ID for any new study. Thus, inspection of the shards by the server’s code allows unique IDs to be generated as long as each ID-minting web service is using a distinct prefix for its IDs. The Open Tree of Life project is minting ID’s using the “ot_” prefix; files that entered the system from a different curation tool, phylografter (Beaulieu *et al*., 2012), have IDs that start with the “pg_” prefix. Note that most of the studies currently in the phylesystem data store were imported from phylografter. Phylografter maintains a list of curators for each study, but the phylesystem data store does not have a detailed record of each edit that was made to each study as a part of curation (because phylografter did not record this information).

The directory structure of each shard is simple: a “study” subdirectory holds a series of subdirectories. Each ID has a numeric suffix. The ID prefix and the last two digits of this numeric suffix are used to create the name for a subdirectory inside the study directory. Inside this directory, each study is in subdirectory with a name that corresponds to the study ID in a file name that corresponds to the ID. So, for example, the phylesystem-api code knows to look for study ot_211 at the path study/ot_11/ot_211/ot_211.json inside the git working directory.

### Mirroring on the server

The server running the phylesystem-api maintains two clones of data repositories. The “working” clone is used to save the updated data (steps 10 and 11 in the workflow). These operations respond to the curator client (in step 13) immediately after these operations complete. To push the data to the GitHub clone of the repository, the server first pulls the master branch from the working repository onto a separate “mirror” clone of the repository. This mirror repository is then locked during the push to GitHub operation (step 14). This architecture keeps the working clone free to save other studies while the high-latency push operation completes using the mirror. The update of the mirror from the working repository is very fast because both are on the same server. Therefore, links to the objects in the git database can be used instead of copy operations.

### otNexSON

The Open Tree of Life project uses three different versions of the otNexSON syntax. Determining which version of otNexSON a particular document is using can be accomplished quickly and reliably by checking a “nexml2json” property in the document. All three versions can be easily interconverted using the peyotl library.

The phylesystem and curator JavaScript applications use versions 1.2 and 1.0 of otNexSON, respectively. Tools that do not require low latency transmission and parsing of the data (e.g. the OTI indexing tool and tools that use the trees in phylesystem for supertree operations) read a direct badgerfish (Badgerfish, 2014) conversion of NeXML which we refer to as otNexSON 0.0 in our documentation. otNexSON version 1 formats are slight tweaks to the badgerfish convention for mapping an XML document to JSON. As with badgerfish, key-value pairs that are attributes in XML are recognized by adding an @ symbol before the key name. NeXML allows for unlimited addition of “meta” elements inside a first-class entity to associate annotations with that entity. The Open Tree of Life project uses a set of these meta annotations to introduce information from curators into the study records. The tags used are described at https://github.com/OpenTreeOfLife/phylesystem-api/wiki/NexSON. In the badgerfish convention these meta elements would be placed as JavaScript objects inside an array associated with the “meta” property name. Adhering to this convention would require any code operating on a JSON version of NeXML to search through (potentially long) lists of meta objects for each piece of annotation. The otNexSON 1.0 and 1.2 syntaxes simply augment the badgerfish convention by using the ∼ character at the front of a property name to indicate that the key-value property corresponds to a “meta” element in NeXML. This ensures smaller file sizes and faster property lookup.

The otNexSON 1.2 syntax reorganizes some of the elements of a NeXML document. In this version of the syntax, the OTUs, trees, nodes, and edges are contained in objects with the IDs of these entities as keys. In a direct badgerfish version of NeXML these entities would be objects in an array, but the order of the objects in these arrays is not useful. For example, reordering the node or edge list has no effect on the conformation of a tree. Use of object IDs as keys make object lookup faster, and allows the JSON to be sorted quickly when being stored in the git repository so that trivial changes (such as changing the order of nodes in a node list) will not result in a difference in the stored version of the study. These formats are documented more fully on the phylesystem-api wiki mentioned above.

The nexson_validation subpackage of the peyotl library is used to ensure that a otNexSON file sent to the phylesystem server is valid. The serialization routines in peyotl also perform re-ordering of the elements within otNexSON files to prevent large commits that would result from, for example, re-ordering the list of OTUs in a file or rotating left / right children of internal nodes. The peyotl library also provides a variety of convenience functions for operating on otNexSON files (including support for exporting the phylogenetic data to widely used formats such as NeXML, NEXUS, and Newick).

### Indexing

A downside of using git as a database is the fact that the use of text files to store data does not inherently provide a way to perform fast lookups of arbitrary objects encoded as text within the files. Performing a text-based search across all studies (i.e. reading through all otNexSON text files) in the phylesystem data store to find all nodes in trees that match a specific set of criteria, for instance, is a prohibitively costly operation to perform on demand. Because the capability to search the phylesystem repository is required for various use-cases (e.g. various aspects of study curation), we implemented an additional tool called OTI (which stands for “open tree indexer”) which parses otNexSON documents and indexes their contents to enable fast searches across studies and their included elements (e.g. trees). OTI is implemented in a Neo4j graph database, and its search features are exposed via publicly available web services. Other Open Tree of Life tools such as phylesystem and the curator application interact with OTI through these web services. Currently available search methods are simplistic, allowing queries based on single properties (e.g. searching for studies by a particular author, or from a particular year) or taxonomic scope (e.g. searching for all trees mapped to any taxon included in some specified higher taxon).

### Webhooks

To create an initial index, OTI reads the entire corpus of studies in the phylesystem data store, but this expensive operation is only required once per deployment of the server. If a study has been added, deleted, or edited by a user and the update was merged to the master branch of the server data store, then an event is triggered (step 12 above) to push the updated data to the GitHub clone of the repository. This assures that the publicly visible version of the data is updated with low latency. In particular, once the push to GitHub has succeeded, a webhook from GitHub posts data about what studies have changed on the GitHub clone to a phylesystem service. This hook triggers the re-indexing of the studies that have changed by the OTI tool so that the searchable cache is kept up to date.

More web services can easily be added to the list of webhooks on GitHub. For example, if someone were to write a service to calculate a statistic on each study in the phylesystem corpus, that programmer could keep the statistics up to date by registering another web hook. The payload of the web hook identifies the files that were changed in a git event. Because the file name portion of each file path corresponds to the study ID, it is trivial for a service receiving the web hook to determine the IDs of the studies that require recalculation.

For services that can tolerate high latency, it is easy to use a scheduled job (e.g. a cron job) to frequently pull the data from the GitHub clone of the repository. Because only the altered studies will have their files touched in the git pull operation, tools such as “make” can be used to update cached calculations for only the studies that have changed.

## 8 Discussion

The phylesystem component of the Open Tree of Life web infrastructure was built to fulfill a critical need for that project: in the process of producing a synthetic estimate of what we know about the phylogeny of life on Earth (Smith *et al*., 2014), information from published studies had to be curated. Phylogenetic hypotheses are simultaneously the products of analyses and the raw material for future analyses. A phylogenetic data store has the advantage of collapsing potentially enormous raw sequence datasets into compact files capturing the inferred evolutionary history of studied taxa (Ané and Sanderson, 2005). However, the raw output of phylogenetic analyses are nearly never sufficient for immediate re-use by other researchers. This curation primarily consists of correcting the rooting of the tree (so that it reflects the finding of the published study), adding metadata about the data or analyses that produced the tree, and mapping the OTU labels to taxa in a taxonomy. Therefore, having a specialty data store to capture the appropriate metadata is essential.

The phylesystem tool was designed to fill this role in way that would make the curated data as widely available as possible. By using git as a primary data store, the system allows other interested parties to easily maintain local clones of all the data.

Data archives in bioinformatics have a wide range of goals and requirements. Using git in place of a typical database is feasible for only a small set of uses which do not require fast processing of a large number of requests. When it is feasible to use distributed version control as a data store, there are several benefits that make this approach appealing. Provenance information in the form of a commit message associated with an identification of the user creating the edit is stored “for free” in such an architecture. Each modification to the data store is backed up efficiently using the VCS’s “push” functionality. The corpus of the data store can be made openly available in a form that is very convenient to other bioinformaticians. Rather than having to unpack a new snapshot and write a script to identify what information has changed since the last snapshot was retrieved, a user can easily pull down the latest changes (with a “git pull origin master” command in the case of a git-based store). Not only will this update be fast, the user knows that he/she can back up to a previous version of the data store if needed using the standard version control features.

The file format of the data to be versioned also has an impact on whether or not it is feasible to use a VCS as a data store. The native tools for comparing versions of a file are line-based. Ideally, such a system would version file formats in which: (a) each datum in a collection is described on a different line, (b) each line is relatively independent, and (c) the order of elements in the serialized file can be made consistent. The otNexSON format that we are currently using is not ideal in these respects. Some operations, such as rerooting a tree, affect many lines in a file (i.e. many branches change source to target orientation). Thus far, the rate of curation has been low enough that most merges have been unambiguous because only one branch of the git history has changed a particular study. The phylesystem currently avoids potentially incompatible merges, and warns the user who committed later that his/her changes have been saved but not merged onto the primary branch. A diff and merge tool that operates on the object model has been written and is currently under testing.

As discussed by others (Drew *et al*., 2013; Magee *et al*., 2014), the rate of deposit of phylogenetic estimates into public archives is currently low. It is also clear that the trees available in digital archives often need some curation. This is a large task because the number of phylogenies published each year is large. Furthermore, some aspects of the curation (most notably verifying that the tip labels are correctly aligned to a taxonomy) require a significant amount of expertise and time investment. It is unclear whether there is a way to motivate the broader community of systematic biologists to invest their time in helping curate a collection of phylogenetic knowledge like phylesystem. Many of the design decisions behind phylesystem reflect a desire to alleviate some potential concerns of data curators. By making the data store publicly accessible as flat files which can be synchronized using robust version control operations, we have tried to lessen concerns that the curation effort is being donated to a resource which might disappear after the end of the Open Tree of Life project. By preserving the history of each commit, we hope to make the data transformation process more transparent, but also make it easier for curators to obtain proper credit for their work.

The design of the data store was also intended to motivate other bioinformaticians to build tools to work with these data. In a traditional database-driven resource the code used to pull information from the private database is quite distinct from the code written by users of the data. However, in phylesystem the server code and client code both deal with the same JSON file format. Thus, developers can easily reuse the code-base of the phylesystem-api as they write new functionalities that use data from phylesystem or even host their own web services using the data. Almost all of the git operations and otNexSON handling operations are implemented in the standalone library, peyotl. This is intended to make it easier for other programmers to clone the data store and work with it locally.

## 9 Conclusion

We have developed a git-based data store for archiving and curating phylogenetic estimates of species relationships. By incorporating curation into the data storage we have lowered the activation cost of entering data into an archive while also allowing continued curation, whether by the original authors or researchers interested in re-using these data, to improve the associated metadata. Using the git VCS allows us to track data curation and maintain provenance, while simultaneously making it straightforward for researchers to maintain their own updatable copies of the database.

## Acknowledgement

We thank members of the Open Tree of Life project for input on design of the system; Rick Ree and Peter Midford for implementing export features for phylografter to allow the phylesystem data store to be populated with studies from phylografter; and the biologists who improve the quality of data by adding and curating studies. Jonathan “Duke” Leto wrote some of the code for a preliminary version of the phylesystem-api tool which served as the initial code base for the current implementation.

**Funding:** We thank NSF AVATOL #1208809, HITS, and an Alexander von Humboldt award to EJM for funding.

